# Failure to invest below-ground may limit the Northern expansion of invasive knotweed: lessons from a two-phase transplant experiment

**DOI:** 10.64898/2026.03.18.712549

**Authors:** Sophie Karrenberg, Elena Barni, Oliver Bossdorf, Hanna Danko, Elisa Giaccone, Madalin Parepa, Christina L. Richards, Nicole Sebesta, Ramona-Elena Irimia

## Abstract

The ecological and evolutionary processes determining species range limits remain poorly understood. Ultimately, range limits depend on the species’ abilities to persist under heterogeneous conditions, by adaptive differentiation and phenotypic plasticity, including transgenerational effects. To investigate ecological differentiation and transgenerational effects in the clonal invasive knotweed, *Reynoutria japonica*, in Europe, we conducted a two-phase transplant experiment: plants sampled along the entire latitudinal gradient were planted in three sites located at the northern range margin, mid-range and near the southern range margin, and then re-transplanted among all three sites after two years. Biomass production and allocation were generally not associated with latitude of origin and previous growth at the same site did not promote performance. We therefore find no evidence that adaptive differentiation or transgenerational effects contribute to the wide distribution of *R. japonica* in Europe. However, at the northern site, with a 25% shorter season, knotweed plants invested much less biomass below-ground, and the pattern was further strengthened in plants that had grown in the northern site in the previous generation. Overwintering below-ground rhizomes are essential for survival and spread. We further explored limiting climate conditions in a species distribution model for the European range and found that mean annual temperature and temperature annual range are the main predictors of the European distribution of *R. japonica*. Taken together, our study suggests that low temperatures and associated short seasons may pose a limit to the broad environmental tolerance of *R. japonica* and restrict its northward spread by reducing below-ground biomass accumulation.

## INTRODUCTION

Processes shaping species distributions and range limits remain poorly understood for most organisms (e.g., Radomski et al., 2026). The capacity to respond to heterogeneous environmental conditions is a key determinant of range size (Colautti et al., 2010; Barton, 2024, Eriksson and Rafajlović, 2022). Whereas adaptation entails heritable differences in trait means or trait plasticities between populations, plastic responses can be reversible in the next generation (Richards et al., 2006; Savolainen et al., 2013). Phenotypic plasticity can occur within and across sexual and clonal generations (transgenerational effects) and can play an important role in range dynamics, and tolerance of temporary adverse conditions, particularly when levels of standing genetic variation for ecologically important traits are limited (McAndry et al., 2025; Latzel and Klimešová, 2010; Mounger et al., 2021). Transgenerational effects are thought to be facilitated by heritable non-genetic effects, such as DNA methylation and histone modification (epigenetic inheritance), as well as by small RNAs, hormones and parental condition (Sommer, 2020; Mounger et al., 2021). A powerful tool for evaluating adaptation and plasticity throughout species ranges are reciprocal transplant experiments under natural or near-natural conditions. Such experiments have offered valuable insights into the prevalence of local adaptation, and the factors driving adaptive differentiation in plant populations (Savolainen et al., 2013; Leimu and Fischer, 2008). While local adaptation is common, even in recently introduced species and in clonal species, it is not universal (Leimu & Fischer 2008; Oduor et al., 2016). In contrast, the role of phenotypic plasticity and transgenerational effects for range dynamics is not well understood. Adaptive transgenerational effects have been demonstrated under field conditions in sexually reproducing plants (e.g., Galloway and Etterson, 2007) and were suggested also for clonally reproducing species (e.g., DuBois et al., 2020). In this study, we use a two-phase transplant experiment to test for both adaptive differentiation and adaptive transgenerational effects in invasive knotweed (*Reynoutria japonica*) and we then related our findings to the limiting climatic conditions identified in a species distribution model.

While fitness in sexually reproducing plants is often assessed using seed output or biomass, performance in clonally reproducing plants is typically estimated by biomass production alone (Younginger et al., 2017). In many non-woody angiosperms, all above-ground biomass dies off during the winter and therefore do not contribute to survival and reproduction in the next season for which storage organs, such as below-ground rhizomes, are crucial. For this reason, investment in below-ground structures is an important performance measure in rhizomatous species (e.g., Xie et al., 2016). In general, higher investment in below-ground and storage organs is found in colder and drier environments and in grasslands, compared to shrublands and forests (Ma et al., 2021; Qi et al., 2019). Below-ground biomass accumulation at Northern latitudes has been identified as an important component of global carbon storage (Ma et al., 2021; Qi et al., 2019). In line with these findings, physiological experiments (reviewed in Poorter et al., 2012) revealed that a reduction in temperature commonly lead to increases in below-ground investment; however, other environmental factors such as light intensity as well as nutrient and water availability also have strong effects on below-ground investment (Poorter et al., 2012; Xie et al., 2016). Population differentiation within species for below-ground investment under controlled conditions was detected in some cases, for example, in silver birch, where populations from colder climates had higher below-ground biomass than those from warmer climates (e.g., Tenkanen et al., 2021). Other species, however, did not show this pattern in experimental settings (D’Hertefeldt et al., 2014; Li et al., 1998) or in field surveys (Chen et al., 2024). Increases in below-ground investment are generally seen as a conservative resource allocation strategy that may be beneficial in challenging environments and can therefore be adaptive in colder climates (e.g., Chapin et al., 1990). However, it is unclear whether biomass allocation to above- and below-ground structures commonly varies within widely distributed species and thereby contributes to range dynamics.

*Reynoutria japonica* Houtt. (Japanese knotweed, Polygonaceae) is a dioecious polyploid plant native to Asia and has a perennial life history (Beerling et al., 1994). The species was introduced to Europe and North America in the past centuries where it has become widespread due to aggressive clonal reproduction with extensive underground rhizomes and is now considered one of the worst invasive species in the world (Lowe et al., 2000; Child and Wade, 2000; Zhang et al., 2024; Lavoie, 2017). Introduced populations of *R. japonica* in both Europe and North America are derived from a single, octoploid lineage originating from Japan (Zhang et al., *2*024). Introduced populations are morphologically similar to putative source populations from Japan but differ from Chinese populations (Cao et al., 2024; Wang et al., 2025). Low temperatures and growing season length have been suggested to determine the Northern range margin of *R. japonica*, whereas summer drought may limit its Southern expansion (Beerling, 1995; Andersen & Elkinton 2023; Zhang et al., 2024); however, information on limiting climatic conditions is incomplete. Transplant experiments have shown pronounced phenotypic plasticity in *R. japonica* and mixed evidence for local adaptation (VanWallendael et al., 2018; Yuan et al., 2024; Cao et al., 2024; Irimia et al., 2025a). Two of these experiments also revealed differences in below-ground investment between transplant sites (Cao et al., 2024; Van Wallendael et al., 2018), but it is unclear how these relate to environmental conditions. Therefore, it remains unresolved whether ecological differentiation or plasticity in biomass allocation patterns contributes to the range dynamics and invasion success of *R. japonica*.

In this study, we investigate mechanisms underlying range limits of *R. japonica* in its European introduced range (Fig. 1). We sampled *R. japonica* populations along a latitudinal gradient and used them in a two-phase transplant (Fig. 2) experiment at three sites across the species latitudinal range. We determined biomass production, as well as above-ground and below-ground biomass allocation, to assess ecological differentiation and transgenerational effects. In addition, we explored limiting climate conditions in a species distribution model for the European introduced range of *R. japonica,* extending previous distribution modeling approaches for this species (Anderson & Elkinton 2023; Zhang et al., 2024). We addressed the following questions: **(1)** Is there ecological differentiation along latitude, or evidence for adaptive differentiation (best performance of plants originating from latitudes similar to the transplant site)? **(2)** Are there transgenerational effects, in particular adaptive transgenerational effects (better performance when transplanted to the same site again)? **(3)** Which climatic conditions limit the European range of *R. japonica*?

**Figure 1.**
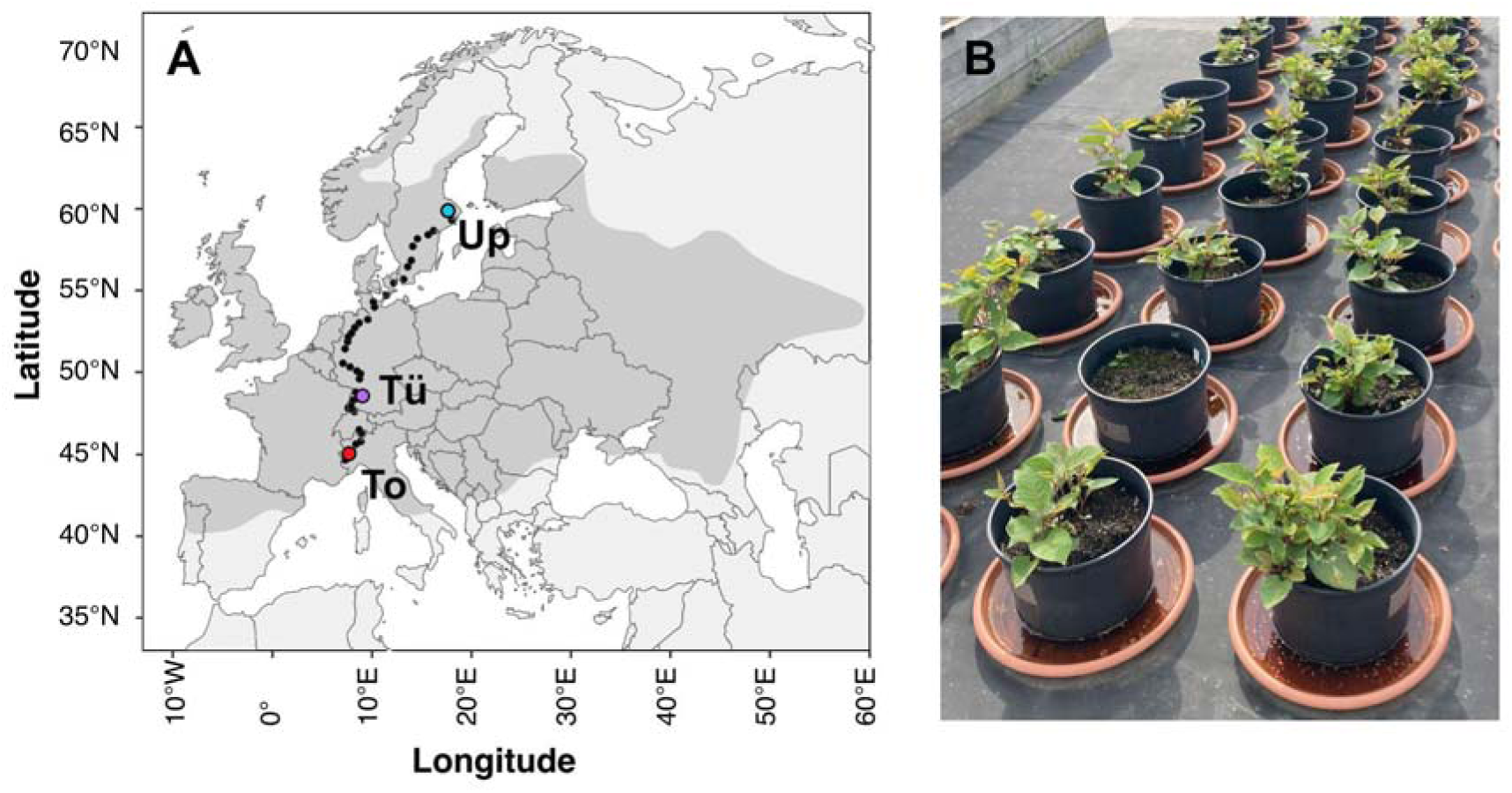
**A.** Range distribution of knotweed, *Reynoutria japonica*, in its introduced range in Europe, based on occurrences reported on GBIF. Black dots indicate 40 sampling locations along a latitudinal gradient and colored dots the location of three transplant sites, Torino (To), Tübingen (Tü) and Uppsala (Up). **B.** Photo of the second phase of the experiment at the Uppsala site.

**Figure 2.**
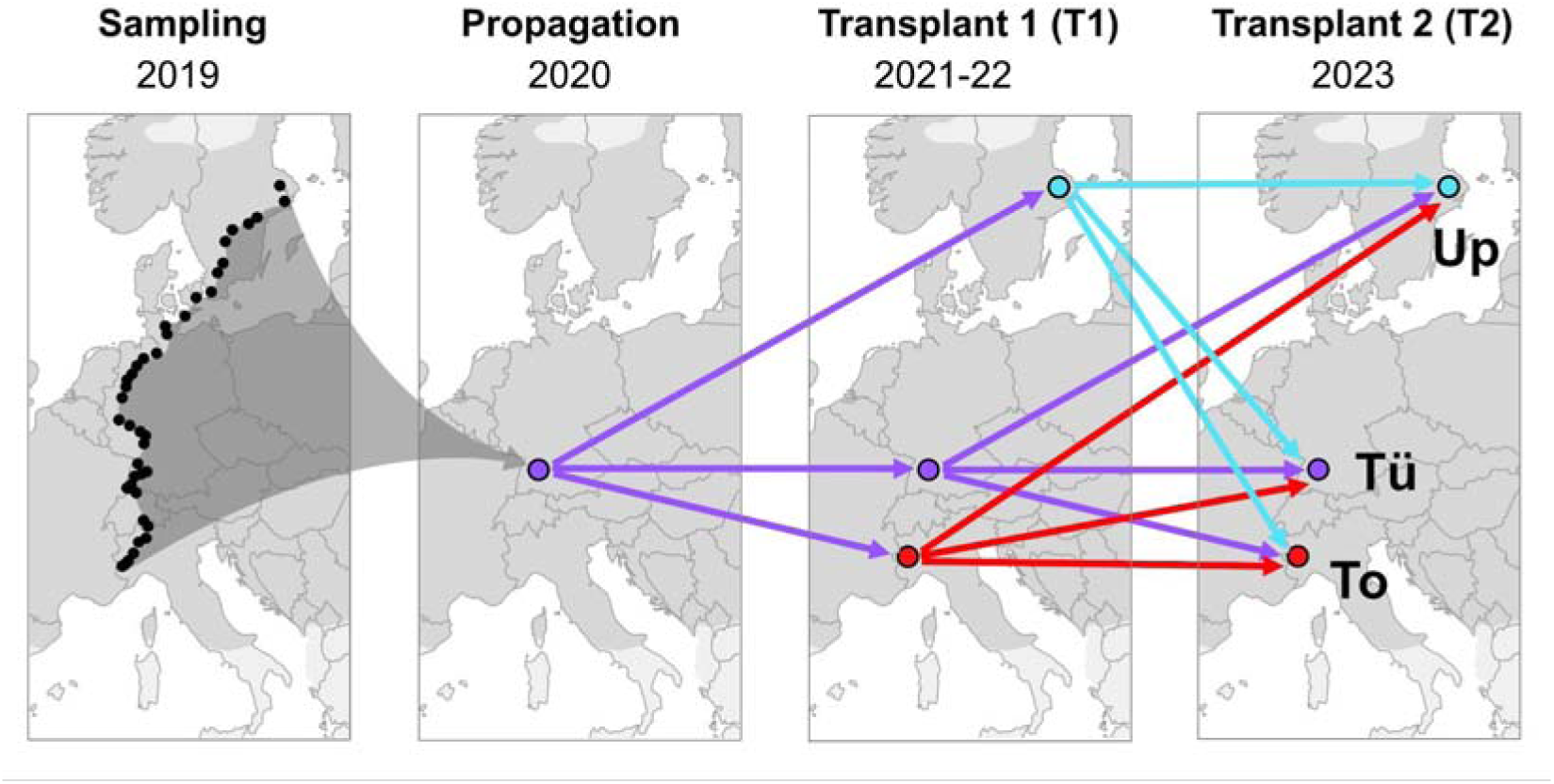
Schematic representation of transplant experiments with the invasive knotweed, *Reynoutria japonica*: sampling from 40 locations along a latitudinal gradient in 2019, propagation at Tübingen University 2020, a first transplant experiment (T1), 2021-2022, to three sites throughout the European invasive range, Uppsala (Up, near Northern range margin), Tübingen (Tü, mid-range) and Torino (To, Southern range margin) and a second transplant experiment (T2) to the same sites that received rhizomes previously propagated at all three transplant sites.

## MATERIAL AND METHODS

### Plant materials and transplant experiments

In summer 2019, we sampled rhizomes from 50 introduced *R. japonica sensu latu* populations along a 2000 km latitudinal transect (44°N, Northern Italy to 59°N, Central Sweden), encompassing a wide climatic gradient and much of the species’ latitudinal range in Europe (Fig. 1). At each site, we excavated five rhizomes along a 30 m transect (see Irimia *et al*., 2025b for more details about the field sampling). We propagated rhizomes in a greenhouse at the University of Tübingen, Germany for two years to reduce environmental effects and harvested three ca. 10 cm long rhizomes from each plant to be used for transplant experiments.

We established three common gardens spread across the European introduced range of *R. japonica*: one in Torino, Italy (45.05°N, 7.68°E, 235 m a.s.l.) near the southern range margin; one in Tübingen, Germany (48.54°N, 9.04°E, 480 m a.s.l.) in the middle of the range and one in Uppsala, Sweden (59.82°N, 17.65°E, 15 m a.s.l.) close to the northern range margin (Irimia et al., 2025b) (Fig. 1).

To investigate ecological differences across the introduced range as well as short-term environmental effects, we carried out a two-phase transplant experiment (Fig. 2). In the first transplant experiment (T1), each common garden received one rhizome cutting from each individual (5×50=250 rhizomes) to be cultivated in a randomized block design with five blocks. We used 25 cm diameter (10 L) pots with commercial potting soil (Patzer Einheitserde Topfsubstrat Type 1; Patzer Erden GmbH, Sinntal, Germany; 340 mg N/L, 260 mg P_2_O_5_/L, 330 mg K_2_O/L, 100 mg Mg/L) and two pellets of slow release fertilizer applied once each year (Osmocote Exact Protect, 5-6M, 14-8-11 + 2MgO + trace elements; Everris International B.V., Heerlen, Netherlands) and watering as needed. After two years (2021 and 2022), we harvested three rhizomes of ca. 10 cm length from three individuals per site of origin, where possible. In the second-phase of the experiment, we redistributed one rhizome cutting from each individual to each of the three gardens for a second transplant experiment for one growth period (2023).

We set up the second transplant experiment (T2) in a randomized block design with three blocks and similar cultivation conditions as the first one, but with 75% soil: 25% vermiculite (Floragard Bio-Erde “Vielseitig” soil composition: 100 mg N/L, 160 mg P_2_O_5_/L and 1100 mg K_2_O/L; and vermiculite: pH value: (CaCl2) 6.0 - 8.0, salt content: (KCl) < 0.5 g/l, conductivity < 200 ms/cm) and liquid fertilizer (Wuxal Top N 12-4-6 (12% N, 4% P2O5, 6% K2O) applied three times, to facilitate the estimation of below-ground biomass. For the second phase of the experiment, we used rhizomes originating from populations from 40 of the 50 sites, excluding four populations later identified as *R.* x *bohemica* and six populations where individuals did not produce a sufficient number of rhizomes for the second transplant experiment. In this study, we report results for this subset comprising three individuals originating from each of 40 sites (Supplementary Material, Table S1, 3 individuals x 40 sites x 3 common gardens; 360 individuals in total).

### Measurements and data analysis

At the end of the first transplant experiment, we determined the fresh mass of each harvested rhizome cutting that was used for the second transplant experiment in 2023, planted February 9 in Uppsala, February 10 in Torino and February 13 in Tübingen. We recorded the emergence of shoots on two dates per site, as well as survival and flowering. We harvested plants after growth cessation due to low temperatures in Torino (October 18) and Tübingen (October 17) and shoot die-off due to frost in Uppsala (October 10). We dried above-ground and below-ground tissues at 60°C for three days and determined dry biomass to the nearest mg.

All statistical analyses were conducted in R version 4.5.1 (R Core Team, 2025). We used linear mixed models in the *lme4* package with a normal distribution of errors (Bates et al., 2015) to analyze how total, above-ground and below-ground biomass as well as the proportion of below-ground biomass were affected by the fresh mass of the rhizome before transplantation (RM), the latitude of origin (Lat) and the first and second transplant gardens (T1 and T2). Models included two-way interactions between the second transplant garden and the other three model effects, rhizome mass, latitude of origin and first transplant garden. Blocks in the second transplant garden were added as a random effect. We used the following model formula, where Y stands for the responses (biomass measures) in the models:

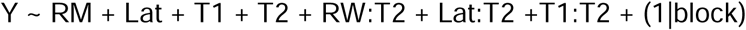

Rhizome mass and latitude of origin were standardized before analysis to allow for interpretation of interaction effects (Forstmeier and Schielzeth, 2011). Responses (biomass measures) were transformed (see Table 1) to improve the distribution of residuals and model fit was assessed by normal quantile-quantile plots and Tukey Anscombe plots.

**Table 1.** Mixed model results for above-ground (**A**), below-ground (**B**) and total biomass (**C**), as well as for the proportion of below-ground biomass (**D**), in a two-phase transplant experiment with the invasive knotweed *Reynoutria japonica* at three sites in Europe, near the southern range margin, mid-range and near the northern range margin. Fixed effect predictors are initial rhizome mass before planting (RM), latitude of origin (Lat.), first transplant site (T1), second transplant site (T2) and two-way interactions between T2 and the other predictors. Blocks in T2 sites were included as a random factor. Response transformations are given as a footnote to each table. Num. df, numerator degrees of freedom; Denom. df, denominator degrees of freedom.

| Predictor | Num. df | Denom. df | F-ratio | P |
| --- | --- | --- | --- | --- |
| **RM** | **1** | **333.61** | **4.434** | **0.0360 \*** |
| Lat. | 1 | 331.99 | 0.195 | 0.6595 |
| **T1** | **2** | **335.89** | **8.042** | **0.0004 \*\*\*** |
| **T2** | **2** | **5.74** | **64.833** | **0.0001 \*\*\*** |
| **RW:T2** | **2** | **333.59** | **6.072** | **0.0026 \*\*** |
| Lat:T2 | 2 | 331.99 | 0.086 | 0.9176 |
| T1:T2 | 4 | 335.49 | 1.403 | 0.2327 |

<sup>a</sup> transformation: $Y^* = \log(Y+20)$

**B. Below-ground biomass<sup>b</sup>**
| Predictor | Num df | Denom df | F | P |
| --- | --- | --- | --- | --- |
| **RM** | **1** | **332.76** | **23.820** | **1.64\*10<sup>-6</sup> \*\*\*** |
| Lat. | 1 | 331.88 | 0.110 | 0.7407 |
| **T1** | **2** | **335.37** | **14.033** | **1.40\*10<sup>-6</sup> \*\*\*** |
| **T2** | **2** | **5.75** | **45.891** | **0.0003 \*\*\*** |
| **RW:T2** | **2** | **332.75** | **3.132** | **0.0449 \*** |
| Lat:T2 | 2 | 331.88 | 0.382 | 0.6829 |
| T1:T2 | 4 | 335.25 | 2.421 | 0.0482 |

<sup>b</sup> transformation: $Y^* = \log(Y+10)$

**C. Total biomass<sup>c</sup>**
| Predictor | Num df | Denom df | F | P |
| --- | --- | --- | --- | --- |
| **RM** | **1** | **332.71** | **18.800** | **1.93\*10<sup>-5</sup> \*\*\*** |
| Lat. | 1 | 331.89 | 0.011 | 0.9150 |
| **T1** | **2** | **335.24** | **12.417** | **6.28\*10<sup>-6</sup> \*\*\*** |
| **T2** | **2** | **5.77** | **26.536** | **0.0012 \*\*** |
| **RW:T2** | **2** | **332.70** | **5.481** | **0.0046 \*\*** |
| Lat:T2 | 2 | 331.89 | 0.248 | 0.7808 |
| T1:T2 | 4 | 335.13 | 2.117 | 0.0783 |

<sup>c</sup> transformation: $Y^* = \log(Y+50)$

**D. Proportion below-ground biomass<sup>d</sup>**
| Predictor | Num df | Denom df | F | P |
| --- | --- | --- | --- | --- |
| **RM** | **1** | **332.84** | **26.964** | **3.62\*10<sup>-7</sup>** *** |
| Lat. | 1 | 331.91 | 1.678 | 0.1961 |
| **T1** | **2** | **335.51** | **6.019** | **0.0027** ** |
| **T2** | **2** | **5.78** | **268.675** | **2.00\*10<sup>-6</sup>** *** |
| **RW:T2** | **2** | **332.83** | **3.047** | **0.0488** * |
| Lat:T2 | 2 | 331.91 | 1.259 | 0.2854 |
| **T1:T2** | **4** | **335.37** | **3.240** | **0.0125** * |

<sup>d</sup> transformation: $Y^* = \text{asin}(Y^{1/2})$

We used the *effects* and *multcomp* packages (Fox and Weisberg, 2019; Hothorn et al., 2008) to perform multiple comparisons (Tukey tests) between transplant gardens for models with significant main effects and to extract model effects, back-transformed to the original scale. Where interaction effects with the second transplant garden were significant, we conducted similar analyses for each garden separately.

### Species distribution modelling

We performed species distribution modeling of *R. japonica* within the European introduced range; this is of interest because the presumably single, clonally reproducing genotype that gave rise to all European introduced populations (Zhang et al., 2024) differs morphologically from other Asian populations (Irimia et al., 2025b; Wang et al., 2025; Cao et al., 2024) and may also have different ecological tolerances. In contrast to previous distribution models of *R. japonica* (e.g., Anderson & Elkinton 2023; Zhang et al., 2024), we used matched occurrences and climate records from a recent period (2000-2016) and uncorrelated climate variables. We constructed the species distribution models (Phillips et al., 2006) based on species occurrence data from the GBIF (GBIF.org) and iNaturalist (https://www.inaturalist.org) databases between 2000 and 2016 and bioclimatic data from CHELSA climate database (https://www.chelsa-climate.org/) for the same period. We included only occurrences that were human observations or preserved specimens (in GBIF data) and had a positional uncertainty < 250 m. To reduce spatial clustering, we thinned all occurrences spatially to the size of a raster cell (2.5 arc min). Finally, we removed coordinates without bioclimatic data. The final dataset included a total of 13,409 species localities across Europe (Supplementary Material, Table S2). For comparison, we also extracted climate data (2000-2016) for occurrences of octoploid *R. japonica* in the putative source region in Japan (Pashley 2003; Ohsawa et al., 2017; Supplementary Material, Table S3).

We calculated 19 bioclimatic variables from monthly minimum and maximum temperature and precipitation data using the *biovars* function implemented in the *dismo* package (Hijmans et al., 2024, Supplementary Material, Table S4). Additionally, we used the first Julian day with a daily mean temperature above 5°C (gdgfgd5) from the CHELSA database, as an indicator of the onset of the growing season (https://www.chelsa-climate.org). We resampled all environmental predictors to a common spatial resolution of 2.5 arc min. To reduce multicollinearity among the variables, we applied the *vifcor* procedure implemented in the *usdm* package (Naimi et al., 2014), which iteratively removes variables based on variance inflation factors until all remaining predictors have correlations below a threshold, here set |r| < 0.7.

We analyzed the climatic space of *R. japonica* in Europe graphically. We performed a Principal components analysis (PCA) of the selected bioclimatic variables (scaled and centered) for all European grid cells at 2.5 arc min resolution and projected European and Japanese occurrences of *R*. *japonica* onto the resulting environmental space.

We fitted a Maxent model using 5,000 background points that were randomly sampled under a spatially structured scheme: 75% of the background points were drawn from within buffers generated by the BackgroundBuffers procedure of the *megaSDM* package (default settings, Shipley et al., 2025), where the buffer radius was defined as twice the 95th percentile of the minimum inter-occurrence distances, and the remaining 25% of background points were sampled from a region extending up to 300 km. To determine the optimal feature classes (fc) and regularisation multiplier (rm), we used the ENMevaluate function from the *ENMeval* package (version 2.0.5.2, Kass et al., 2021). We varied regularisation multipliers from 1 to 5, in increments of 1, and included all feature classes (L = linear, Q = quadratic, H = hinge, P = product and T = threshold) and their combinations (Muscarella et al., 2014). For cross-validation, we employed a hierarchical spatial checkerboard partitioning (Muscarella et al., 2014). The final model was selected based on the maximal average validation area under the curve, indicating the best combination of feature and regularisation multiplier. We identified climate variables associated with range limits using the prediction importance and a permutation test (Smith and Santos, 2020). For the most influential variables, we defined species climatic limits as the 2.5^th^ and 97.5^th^ percentiles of their values extracted from grid cells with predicted probability of occurrence more than 0.5, thereby approximating their lower and upper bounds.

## RESULTS

### Survival, flowering and growth season length

Almost all of the 120 rhizomes planted at each site produced above-ground biomass (data available as Supplementary Material, Table S5). In Tübingen, all rhizomes survived, whereas 117 survived in Torino and 116 in Uppsala (Fig. 3). Flowering initiation differed strongly between sites: 76 plants flowered in Torino (65.0% of the surviving plants), 37 in Tübingen (30.8%) and only 2 in Uppsala (1.7%).

**Figure 3.**
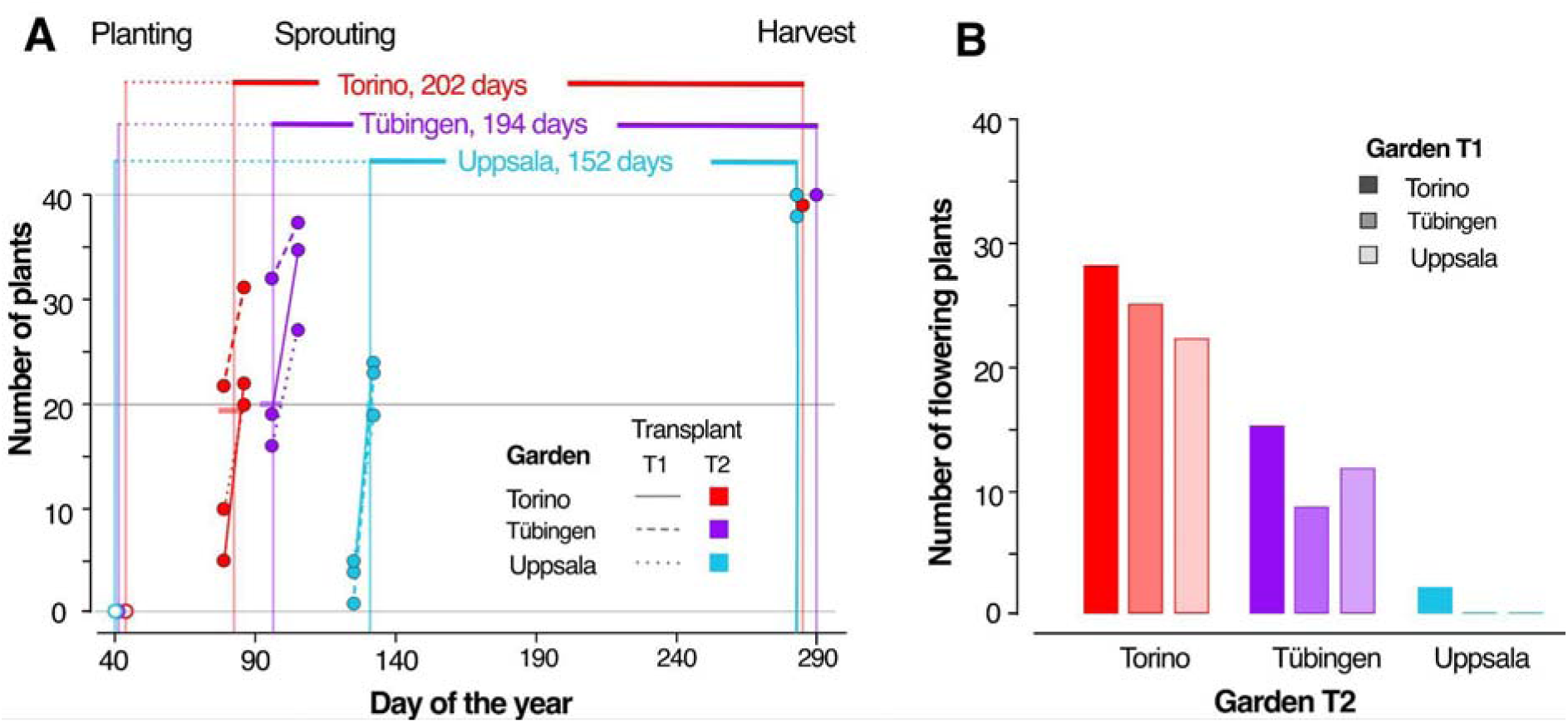
Season length **(a)** and number of flowering individuals **(b)** in a transplant experiment (second transplant experiment, T2, see methods and Figure 2) with the invasive knotweed, *Reynoutria japonica* at three sites, Torino (red), Tübingen (purple) and Uppsala (petrol). Given are numbers of plants with above-ground biomass at two time points during sprouting and at the start and end of the T2 experiment (**A**) or with flowers (**B**). Plants were grown from rhizomes previously transplanted to the same sites (T1, see legends). Season lengths at each T2 transplant site (A) were estimated using the approximate time at which 50% of the rhizomes had sprouted (colored bar and vertical line).

Set up and harvest of the experiment were done at a similar time at all sites in February and October, respectively, but sprouting was observed much later in Uppsala than in Tübingen and Torino (Fig. 3). The approximate date at which 50% of the surviving plants had sprouted was 24 March (day 83) in Torino, 4 April (day 96) in Tübingen and May 11 (day 131) in Uppsala, such that we estimated the median length of the growing period until harvest as 202, 194 and 152 days in Torino, Tübingen and Uppsala, respectively (Fig. 3).

### Current and previous transplant sites, but not latitude of origin, affect biomass

We detected clear evidence for effects of the current (T2) and previous (T1) transplant site on all parameters measured (Table 1, Fig. 4, data available as Supplementary Material, Table S5). In addition, interaction effects between T1 and T2 affected below-ground biomass and the proportion of below-ground biomass (Table 1, see Supplementary Material, Table S6, for results of sub-models within T2 sites). We did not find evidence for effects of latitude and the interaction between latitude and T2 on any of the biomass measures (Table 1).

**Figure 4.**
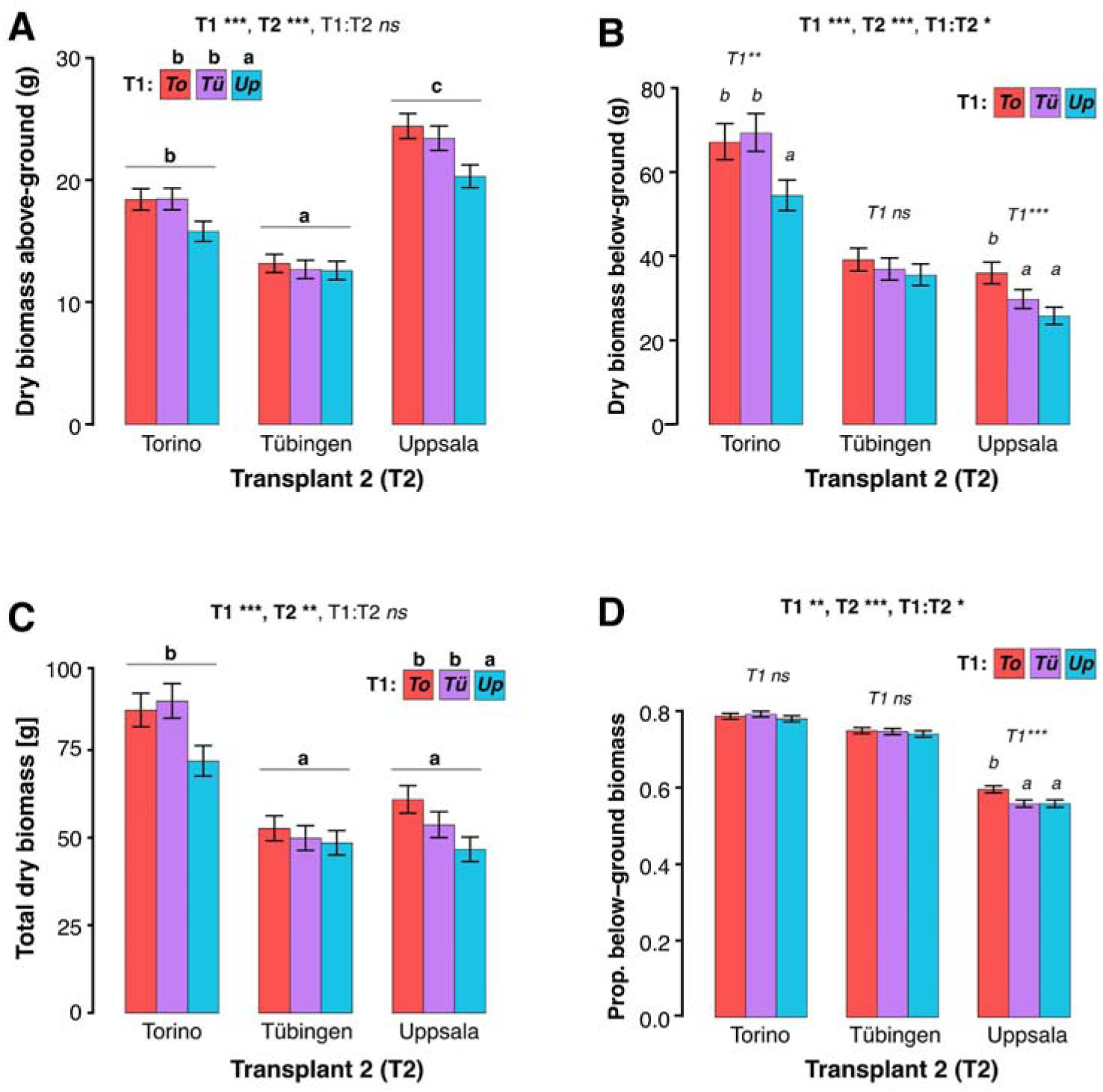
Above ground **(A)**, below-ground **(B)** and total **(C)** dry biomass, as well as the proportion of below ground biomass **(D)** produced by rhizomes of the invasive knotweed *Reynoutria japonica* in a transplant experiment (T2, see Fig. 2) to three European sites, Torino (To), Tübingen (Tü) and Uppsala (Up) after previous growth at the same three sites (T1, To, red; Tü, purple; Up, petrol). Given are predicted marginal effects with one standard error, estimated from mixed models with results of the fixed effects of T1 and T2 sites, as well as the interaction between these factors (T1:T2) above each panel (Table 1). For models without significant interaction effects (A, C), different letters (bold) indicate significant differences between T2 sites (above groups of bars) and T1 sites (above legend). For models with significant interaction effects (B, D), results of T1 effects in sub-models for each T2 site are given in italics (see Supplementary Material, Table S6), together with multiple comparisons among T1 sites where significant T1 effects were detected (italic letters above individual bars). All models also included fixed effects of initial rhizome mass (RM), the interaction between RM and T2 and sampling latitude, as well as transplant blocks as a random effect. ***, P < 0.001; **, P < 0.01; *, P < 0.05; ns, not significant.

We found strong and opposing biomass patterns in the three T2 sites. Above-ground biomass was higher at the northernmost site, Uppsala, than at Tübingen (76% increase) and Torino (29% increase). Below-ground biomass, as well as the proportion of below-ground biomass, in contrast, were reduced at this site (Fig. 4). Below-ground biomass was 52% reduced when comparing Torino plants to those at Uppsala, and 17% when comparing Tübingen plants to Uppsala plants. This resulted in 25% and 21% reductions in the proportion of below-ground biomass for Torino vs. Uppsala and Tübingen vs. Uppsala (Fig. 4). Total biomass, however, was clearly highest at the southernmost site, Torino. Knotweed plants at the mid-range site (Tübingen) produced a total biomass similar to those at the Uppsala site but with a below-ground proportion similar to those at the Torino site (Fig. 4).

The T1 site Torino had positive transgenerational effects on biomass production whereas the reverse was true for plants previously grown at Uppsala. Above-ground biomass and total biomass were 12% respectively 10% reduced when plants were previously grown in Uppsala, as compared to Torino and Tübingen (Fig. 4). At the T2 site Uppsala, below-ground biomass and the proportion of below-ground biomass were 29% and 6% lower, respectively, when plants were previously grown (T1) at the same site, compared to previous growth at Torino (Fig. 4, Supplementary Material, Table S6). At the T2 site Torino, previous growth at the T1 site Uppsala also reduced below-ground biomass by 23% as compared to previous growth at Torino. Previous growth at Tübingen (T1) exhibited similar patterns to either Uppsala-grown plants (at the Uppsala T2 site) or to Torino-grown plants (at the Torino T2 site).

### Stronger increases of biomass with rhizome mass under warmer conditions

The effects of initial rhizome mass (at planting) on all biomass measures differed between the T2 sites, as indicated by significant interaction effects between rhizome mass and T2 (Table 1). At the Torino site, above-ground, below-ground and total biomass strongly increased with rhizome mass, whereas the proportion of below-ground biomass increased only weakly (Fig. 5, Supplementary Material, Table S6). At the Uppsala site, in contrast, higher rhizome mass did not affect above-ground or total biomass but led to a significant increase in below-ground biomass and in the proportion of below-ground biomass (Fig. 5). At the Tübingen site, we did not find evidence for effects of rhizome mass on any of the biomass measures (Fig. 5).

**Figure 5.**
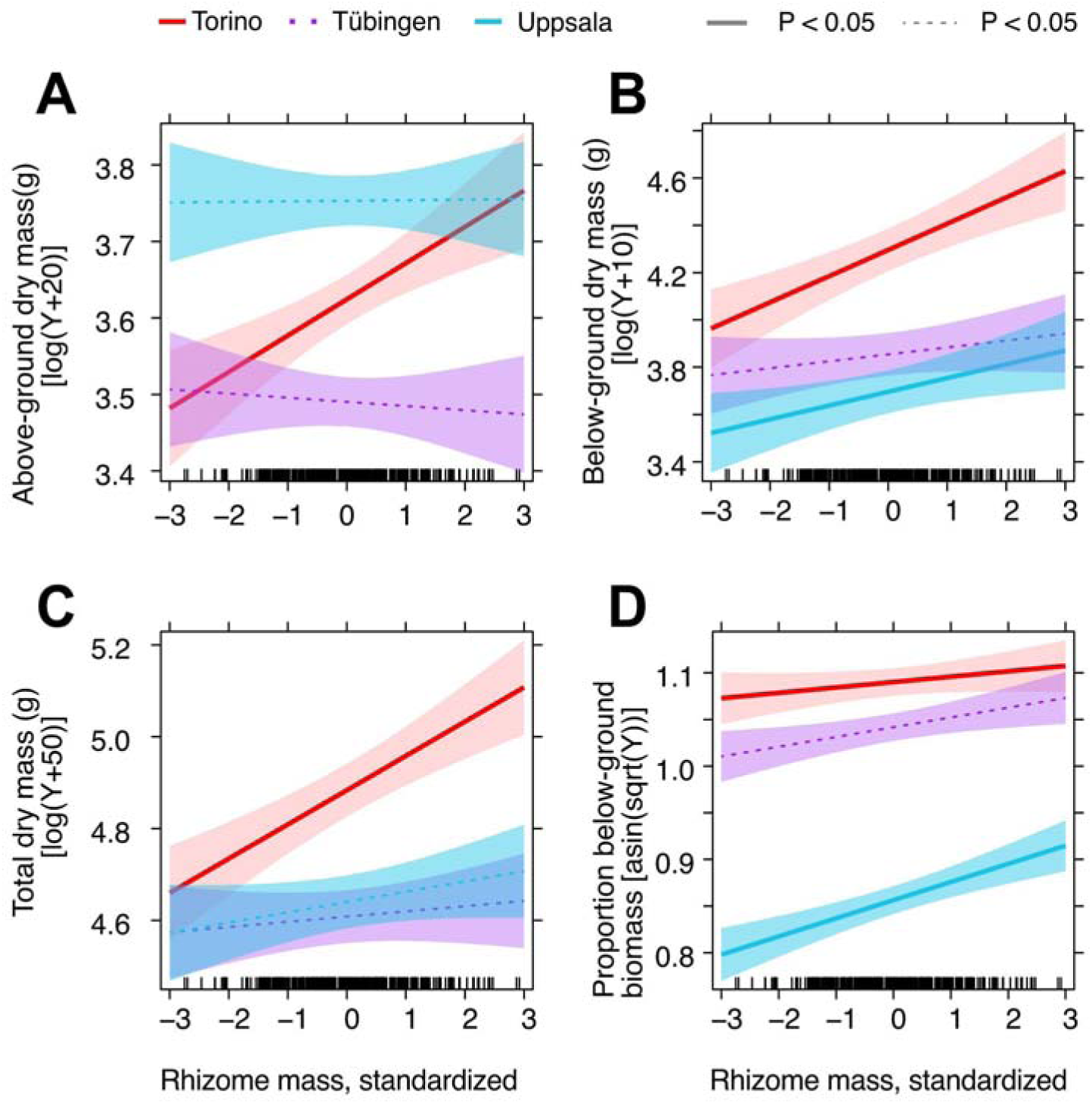
Relationship between the fresh rhizome mass (before planting) to above-ground (A), below-ground (B) and total (C) dry biomass, as well as the proportion of below-ground biomass (D) produced by the invasive knotweed *Reynoutria japonica* in a transplant experiment to three sites in Europe (T2, see Fig. 2), Torino (red), Tübingen (purple) and Uppsala (petrol). Given are predicted effects with 95% confidence limits as estimated from mixed models within each transplant site (Table 1, Supplementary Material, Table S6). Regressions that are significant at *α* = 0.05 are displayed as thick lines and non-significant ones as dashed lines. Ticks above the x-axis show values in the dataset.

### Correlations among climatic variables

In the set of occurrence and background data points used for distribution modelling, many bioclimatic variables were highly correlated (Fig. 6A, Supplementary Material, Fig. S1). A group of temperature variables (bio1, bio3, bio9, bio6, bio11) and the gdgfgd5 (growing season length) formed one correlated group, perpendicular to a precipitation-related group of correlated variables (bio12, bio13, bio14, bio16, bio17, bio19) (Fig. 6A). Further, smaller clusters were formed by bio7 (temperature annual range) and bio4 (temperature seasonality), as well as by bio10 (mean temperature of warmest quarter), bio5 (mean daily maximum near-surface air temperature of the warmest month) and bio2 (mean diurnal near-surface air temperature range), whereas bio8 (mean near-surface air temperature of wettest quarter), bio18 (precipitation of warmest quarter), bio15 (precipitation seasonality) were more independent. Using *vifcor* procedure implemented in the *usdm* package (Naimi et al., 2014), we identified the following set of eight variables with correlations of |r| < 0.7 that we later used for distribution modeling (Fig. 6A, Supplementary Material, Fig. S1): mean annual temperature (bio 1), isothermality (bio3), temperature annual range (bio7), mean temperature of wettest (bio 8) and driest (bio 9) quarter, precipitation of driest month (bio 14) and of warmest quarter (bio18) and precipitation seasonality (bio15).

**Figure 6.**
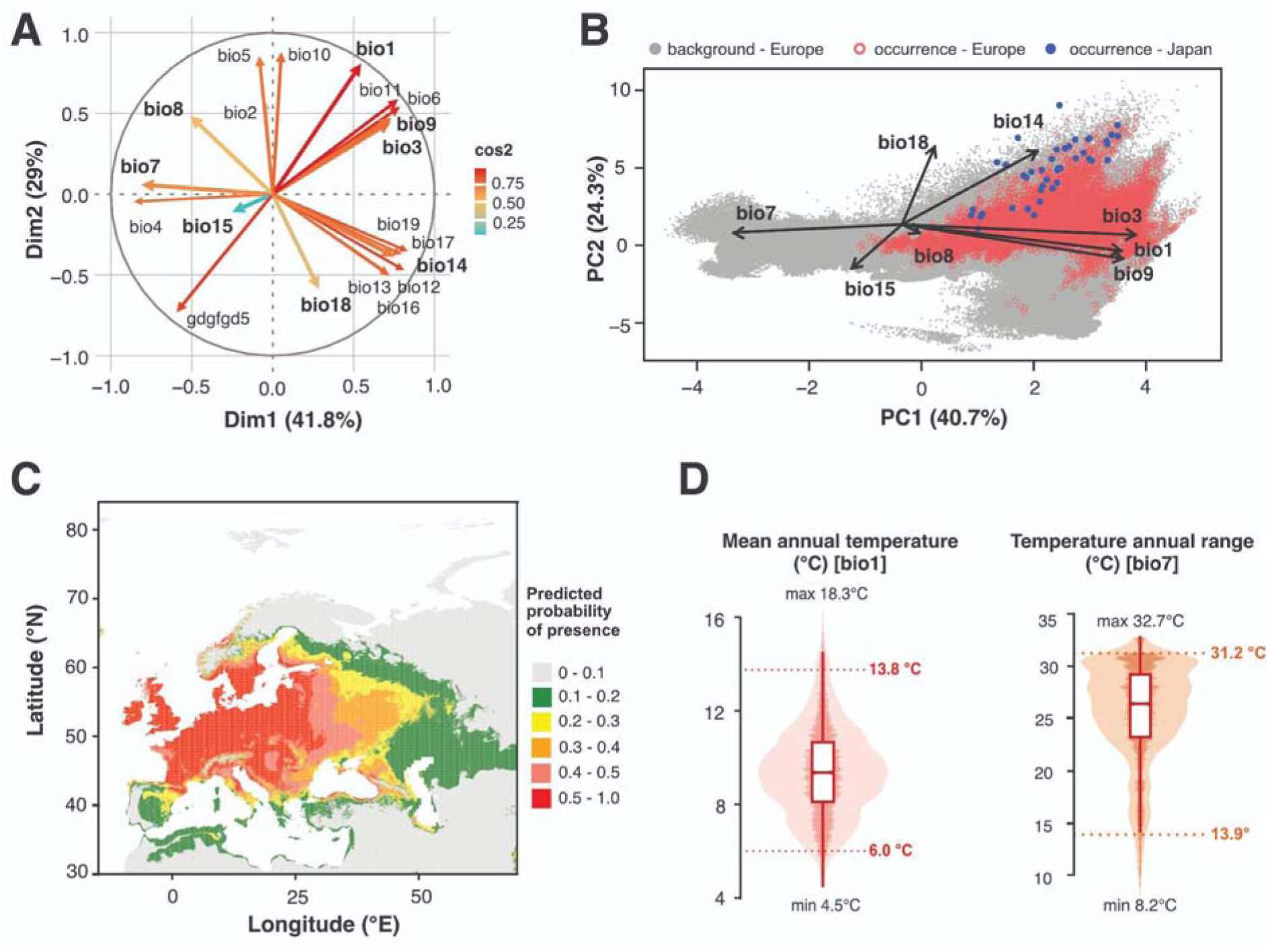
Species distribution model (SDM) for invasive knotweed, *Reynoutria japonica* in its European introduced range based on occurrence and climate data for 2000-2016: **(A)** correlation biplot among bioclimatic variables (occurrences and background), cos2 indicates a representation of the variable on the principal component, variables with |r| < 0.7 are in bold; **(B)** climatic space of *R. japonica* in Europe: Principal Components Analysis (PCA) of European grid cells (2.5 arc, grey) for the selected bioclimatic variables (arrows) with projected occurrences of *R. japonica* in Europe (red) and in the putative source regions in Japan (blue). **(C)** predicted probability of presence in Europe (Maxent model) **(D)** distributions and boxplots of of the main predictors, mean annual temperature (left) and annual temperature range (right), in grid cells with predicted probability of occurrence > 0.5, with climatic limits indicated as 2.5^th^ and 97.5^th^ quantiles (dotted lines).

### Putative source sites are marginal to the climatic space in the introduced European range

The climatic space of *R. japonica* in Europe was projected onto warmer temperatures (bio1, bio3, bio9), smaller temperature annual ranges (bio7) and higher precipitation in the warmest quarter (bio18, Fig. 6B). Projected occurrences of octoploid *R. japonica* in the putative source region in Japan were situated at the margin of the European climate space of *R. japonica,* with higher precipitation in the warmest quarter, as expected for a monsoon climate (bio18, Fig. 6B). Note that these comparisons are difficult to evaluate, because both tetraploid and octoploid populations of *R. japonica* occur in Japan and it is unclear whether they differ ecologically (Pashley 2003; Ohsawa et al., 2017).

### Mean annual temperature and annual temperature range define the distribution

Our distribution model predicted areas of higher suitability for *R. japonica* mostly in the central and northwestern part of Europe, according to the Maxent models (Fig. 6C), in strong agreement with the observed distribution (Fig. 1A). We evaluated Maxent models by comparing AUC (Area Under the Receiver Operating Characteristic Curve) values (Phillips et al., 2006). Training AUC values ranged from 0.73 to 0.82, whereas test AUC values ranged from 0.66 to 0.72, indicating moderate but acceptable discrimination ability for presence-only models. Such AUC values are expected under spatially independent cross-validation and large geographic extents, where environmental overlap between training and test data is limited. The omission rates of test samples at the 10% training presence threshold ranged from 0.10 to 0.11, indicating good model calibration. The optimal Maxent model (selected by highest AUC) used LQHPT (Linear, Quadratic, Hinge, Product, and Threshold features) features and a regularization multiplier of 1. The model showed high predictive performance with a mean test AUC of 0.79 (SD = 0.01) across four spatial cross-validation folds, indicating good discrimination between presence and background locations. The training AUC was 0.82, and the test omission rate at the 10-percentile training presence threshold was 0.11, suggesting minimal overfitting.

Our models had high Continuous Boyce Index (CBI) values (>0.99), indicating a strong discriminatory power in correctly ranking bioclimatic predictors, supporting the robustness of the predictions despite moderate AUC values (Whitford et al., 2024). Among the eight uncorrelated bioclimatic variables retained in the model (Fig. 6A, Supplementary Material, Fig. S1), annual mean temperature (bio1) clearly had the highest predictor contribution and permutation importance (Table 2). The second most important predictor variable was temperature annual range (bio7) whereas the remaining variables had minor contributions to the model (Table 2).

**Table 2.**
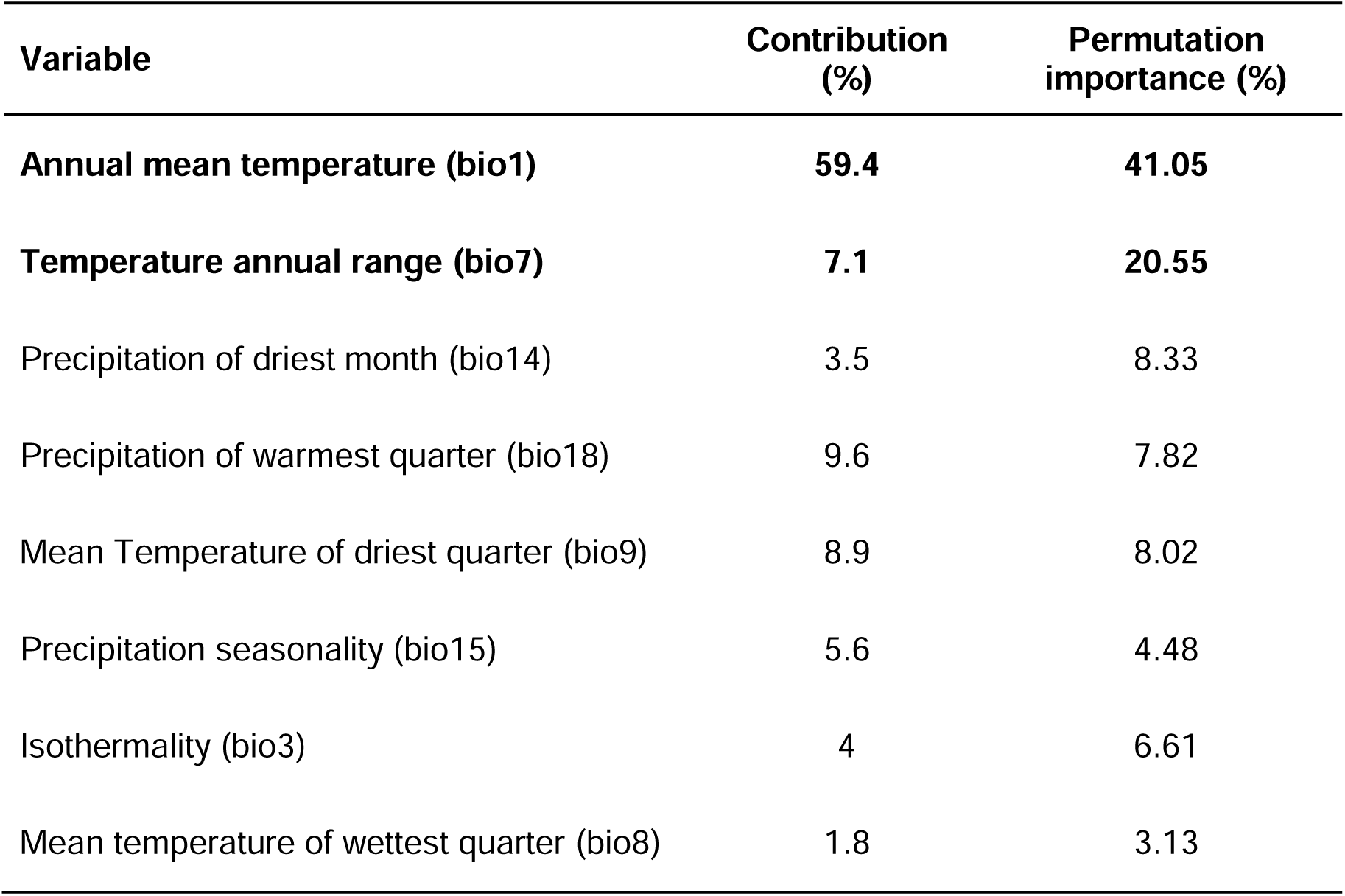
Relative importance of individual bioclimatic variables based on permutation importance from Maxent models for invasive knotweed (*Reynoutria japonica*) in the introduced range in Europe. Variables contributing most strongly to climatic suitability are highlighted in bold. For explanations of bioclimatic variables see Supplementary Material, Table S4.

Our distribution model further predicts that *R. japonica* is expected to occur (with > 50% prediction probability) where the mean annual temperature is between 6.0 and 13.8 °C (2.5^th^ and 97.5^th^ quantiles, minimum: 4.5°C, maximum: 18.3°C) and temperature annual range is between 13.9 and 31.2 °C (2.5^th^ and 97.5^th^ quantiles, minimum: 8.2°C, maximum: 32.7°C, Fig. 6D). It is, however, generally difficult to interpret limits predicted using species distribution models, due to the dependence of these models on the climatic variation in both background and occurrence data.

## DISCUSSION

Our two-phase transplant experiment does not support a contribution of adaptive differentiation or beneficial transgenerational effects to the range dynamics of invasive knotweed *R. japonica* in its introduced range in Europe. Instead, we show that European introduced knotweed reduces below-ground investment when transplanted to near the northern range margin, even though total biomass production is similar to the mid-range site. Interestingly, this pattern is further strengthened by transgenerational effects of the previous year’s conditions, suggesting persistent physiological effects of growth under marginal conditions. According to distribution models, mean annual temperature is a main determinant of the *R. japonica* distribution in Europe, with predicted occurrence at sites with mean annual temperatures between 6.0 and 13.8°C. Below we discuss the role of temperature and associated climatic parameters for reduced below-ground investment and the establishment of northern range limits.

We transplanted rhizomes originally sampled along a latitudinal gradient to three sites spanning the introduced European range and did not detect any effect of latitude of origin on biomass production or allocation. This is in line with previous findings from the first phase of the experiment, in which plant height and other traits were not associated with latitude of origin (Irimia et al., 2025a). Even though we detected transgenerational effects in all biomass parameters measured (see below for a detailed discussion), there was no evidence that plants benefit from transplantation to the same site. These results do not provide evidence for ecological differentiation along latitude or adaptive transgenerational effects. We used one individual from each of 40 different populations per transplant site and cannot make inferences about differentiation between particular population pairs. Instead, we tested whether adaptive differentiation along latitude was detectable as higher performance in populations originating from latitudes similar to the transplant site, a powerful and commonly used approach (Gorton et al., 2022). Note however, that in our study, almost all plants survived, and our conclusions are therefore based on biomass production only. Our results agree with reports from the American introduced range of *R. japonica* where evidence for local adaptation across the range and to different local habitats was mixed (VanWallendael et al., 2018; Yuan et al., 2024). The apparent lack of strong local adaptation in the introduced ranges of *R. japonica* is likely due to clonal reproduction and the origin of the introduced populations from a single lineage (Beerling et al., 1995; Zhang et al., 2024). A lack of local adaptation was also suggested for the clonal invasive species alligator weed, *Alternanthera philoxeroides* (Geng et al., 2007) and the ornamental shrub *Buddleja davidii* (Ebeling et al., 2011). In other invasive species with similar life histories, in contrast, local adaptation was detected, for example in monkeyflowers, *Mimulus spp.,* and cheatgrass, *Bromus tectorum* (Simón-Porcar et al., 2021; Gamba et al., 2025). Overall, our data suggest that local adaptation and adaptive transgenerational effects do not determine the wide distribution of *R. japonica* in its European introduced range. Instead, we provide further support for broad environmental tolerance as the main mechanisms facilitating the spread of this species (Wang et al. 2025, Irimia et al., 2025a).

In our experiment, *R. japonica* strongly reduced below-ground investment at the site near the northern range margin compared to the two other sites. Interestingly, this pattern was further strengthened by transgenerational effects: previous growth at the northern site led to reduced total biomass at all sites and reduced below-ground investment at the northern site. At the same time, plants at the northern site produced the highest above-ground biomass of all sites and a similar total biomass than plants at the mid-range site. A possible explanation for these patterns of biomass accumulation is a mismatch between growth regulation and the 20-25% shorter season at the northern site, as compared to the other two sites (Fig. 3A). Below-ground growth continues later into the season than above-ground growth in many species (Abramoff and Finzi, 2015) and this has also been reported for *R. japonica* (Price et al., 2001). Translocation of assimilated carbon to rhizomes is intensified at the end of the season in *R. japonica* likely including both structural and non-structural (storage) carbohydrates (Price et al., 2001). Insufficient accumulation of storage carbohydrates or other compounds possibly explains the productivity-reducing, transgenerational effects of the northern site, despite a similar mass of transplanted rhizomes from all sites. Shorter seasons have also been suggested to limit the distribution of *R. japonica* at higher elevations in the native range (Maruta 1983). Our results are similar to those obtained under controlled conditions in the rhizomatous weed *Aegopodium podagaria,* where plants at the northern site had higher above-ground and total biomass and reduced below-ground investment (D’Hertefeldt et al., 2014). Previous transplant experiments with *R. japonica* have shown both increased (VanWallendael et al., 2018) and reduced (Cao et al., 2024) below-ground investment at northern sites. However, these studies are difficult to compare to ours because VanWallendael et al. (2018) transplanted plants into forest habitats in North America and Cao et al. (2024) conducted their experiment in more southern sites in China (below 34°N). Reduced below-ground investment close to the northern range margin in Sweden (59°N), as reported here, is detrimental for spread in the next generation and may therefore limit the northward expansion of *R. japonica*.

Our distribution model for the European invasive range of *R. japonica* agrees with the models of Andersen and Elkinton for Europe (2023) and the world-wide model of Zhang et al. (2024). We provide an extended analysis using uncorrelated variables and matched occurrence and climate records. Mean annual temperature (bio1) clearly had the strongest impact on the distribution of *R. japonica* in Europe, followed by temperature annual range (bio7). These variables are associated with the classic plant ecological gradients of temperature and continentality (Bartelheimer and Poschlod, 2016) and are consistent with suggestions of Beerling et al. (1995). In our dataset, mean annual temperature was strongly correlated with season length (Fig. 6A, Supplementary Material, Fig. S1). Our distribution models are therefore consistent with an important role for season length for defining the northern range margin. Temperature annual range is the difference between maximum and minimum annual temperature (daily values averaged per month) and is highly correlated with these variables. Beerling et al. (1995) and Maruta et al. (1983) suggested that tolerance to low temperatures may limit the distribution of *R. japonica*. Indeed, leaves of *R. japonica* are susceptible to frost damage, even by shorter exposure (Baxendale and Tessier, 2015) and early autumn night frosts may also limit the season length, as we have also observed in our experiment. However, the freezing tolerance of overwintering rhizomes under natural conditions remains unknown.

Overall, we found no evidence for differentiation along latitude within the introduced range or for adaptive transgenerational effects. We conclude that these processes may not underlie the invasion success and wide distribution of *R. japonica* in Europe. Instead, broad environmental tolerance is likely the main mechanism facilitating the spread of this invasive species. However, our experimental data and distribution models also suggest that low mean annual temperatures and associated short seasons may constitute a limit to the environmental tolerance of *R. japonica*: northern conditions led to high above-ground productivity but reduced accumulation of below-ground biomass, that was reinforced by negative transgenerational effects from previous growth at the northern site. This suggests that failure to accumulate sufficient below-ground biomass could limit further northward spread on the introduced populations. Our study thus contributes to the understanding of evolutionary and ecological processes that mediate species range limits.

## DATA AVAILABILITY

The raw data for this study is available as Supplementary Material and the R code used for analyses will be deposited on github following manuscript acceptance.

## ACKNOWLEDGMENTS

We thank Christiane Karasch-Whittmann, Sabine Silberhorn, Anaëlle Regen and Mattias Vass for help with the field experiment and processing plant samples in the lab. This study was supported by the German Federal Ministry of Education and Research (BMBF; MOPGA Project 306055 to Christina L. Richards), the German Research Foundation (DFG; grant 431595342 to Oliver Bossdorf and Christina L. Richards) the European Union’s Horizon 2020 research and innovation program under the Marie Skłodowska-Curie grant agreement No 101033168 (to Ramona E. Irimia), and MSCA4Ukraine program of the Humboldt foundation (doctoral fellowship No. 1233613 to Hanna Danko).

## REFERENCES

Abramoff RZ, Finzi AC. 2015. Are above- and below-ground phenology in sync? The New Phytologist 205: 1054–1061.

Andersen JC, Elkinton JS. 2023. Climate suitability analyses compare the distributions of invasive knotweeds in Europe and North America with the source localities of their introduced biological control agents. Ecology and Evolution 13: e10494.

Bartelheimer M, Poschlod P. 2016. Functional characterizations of Ellenberg indicator values – a review on ecophysiological determinants. Functional Ecology 30: 506–516.

Barton N. 2024. Limits to species’ range: the tension between local and global adaptation. Journal of Evolutionary Biology 37: 605–615.

Bates D, Mächler M, Bolker B, Walker S. 2015. Fitting Linear Mixed-Effects Models Using lme4. Journal of Statistical Software 67: 1–48.

Baxendale VJ, Tessier JT. 2015. Duration of freezing necessary to damage the leaves of *Fallopia japonica* (Houtt.) Ronse Decraene: freezing of knotweed leaves. Plant Species Biology 30: 279–284.

Beerling DJ, Bailey JP, Conolly AP. 1994. *Fallopia Japonica* (Houtt.) Ronse Decraene. The Journal of Ecology 82: 959–979.

Beerling DJ, Huntley B, Bailey JP. 1995. Climate and the distribution of *Fallopia japonica*: use of an introduced species to test the predictive capacity of response surfaces. Journal of Vegetation Science 6: 269–282.

Cao P-P, Yin W-D, Bi J-W, Lin T-T, Wang S-Y, Zhou H, Liao Z-Y, Zhang L, Parepa M, Ju R-T, Ding J-Q, Nie M, Bossdorf O, Richards CL, Wu J-H, Li B. 2024. Clonal plasticity and trait stability facilitate knotweed invasion in Europe. Journal of Plant Ecology 17: rtae067.

Chapin FS III, Schulze E, Mooney HA. 1990. The ecology and economics of storage in plants. Annual Review of Ecology and Systematics 21: 423–447.

Chen Y, Wu F, Wang Y, Guo Y, Kirwan ML, Liu W and Zhang Y. 2024. Latitudinal trends in the biomass allocation of invasive *Spartina alterniflora*: implications for salt marsh adaptation to climate warming. Frontiers in Marine Science 11:1510854.

Child L, Wade M. 2000. The Japanese Knotweed Manual. Packard Publishing Limited, Chichester. 123 pp.

Colautti RI, Eckert C, Barrett SCH. 2010. Evolutionary constraints on adaptive evolution during range expansion in an invasive plant. Proceedings of the Royal Society B: Biological Sciences 277: 1799–1806.

D′Hertefeldt T, Eneström JM, Pettersson LB. 2014. Geographic and habitat origin influence biomass production and storage translocation in the clonal plant *Aegopodium podagraria*. PLoS One 9: e85407.

DuBois K, Williams SL, Stachowicz JJ. 2020. Previous exposure mediates the response of eelgrass to future warming via clonal transgenerational plasticity. Ecology 101: e03169.

Ebeling SK, Stocklin J, Hensen I, Auge H. 2011. Multiple common garden experiments suggest lack of local adaptation in an invasive ornamental plant. Journal of Plant Ecology 4: 209–220.

Eriksson M, Rafajlović M. 2022. The role of phenotypic plasticity in the establishment of range margins. Philosophical Transactions of the Royal Society of London. Series B, Biological Sciences 377: 20210012.

Forstmeier W, Schielzeth H. 2011. Cryptic multiple hypotheses testing in linear models: overestimated effect sizes and the winner’s curse. Behavioral Ecology and Sociobiology 65: 47–55.

Fox J, Weisberg S. 2019. An R Companion to Applied Regression. Thousand Oaks, CA, USA.

Galloway LF, Etterson JR. 2007. Transgenerational plasticity is adaptive in the wild. Science 318: 1134–1136.

Gamba D, Vahsen ML, Maxwell TM, Pirtel N, Romero S, Van Ee JJ, Penn A, Das A, Ben-Zeev R, Baughman O, et al. 2025. Local adaptation to climate has facilitated the global invasion of cheatgrass. Nature Communications 16: 10203.

Geng Y-P, Pan X-Y, Xu C-Y, Zhang W-J, Li B, Chen J-K, Lu B-R, Song Z-P. 2007. Phenotypic plasticity rather than locally adapted ecotypes allows the invasive alligator weed to colonize a wide range of habitats. Biological Invasions 9: 245–256.

Gorton AJ, Benning JW, Tiffin P, Moeller DA. 2022. The spatial scale of adaptation in a native annual plant and its implications for responses to climate change. Evolution 76: 2916–2929.

Hijmans R, Phillips S, Leathwick J, Elith J. 2024. dismo: Species Distribution Modeling. R package version 1.3-16. https://cran.r-project.org/package=dismo.

Hothorn T, Bretz F, Westfall P. 2008. Simultaneous inference in general parametric models. Biometrical Journal 50: 346–363.

Irimia RE, Parepa M, Sebesta N, Barni E, Giaccone E, Guo Y, Karrenberg S, Richards C, Bossdorf O. 2025a. Phenotypic plasticity of invasive knotweed across Europe: a distributed common garden experiment. bioRxiv. 10.1101/2025.08.18.667133.

Irimia RE, Zhao W, Cao P, Parepa M, Liao Z-Y, Wang S, Mounger JM, Richardson C, Elkott F, Zhuang X, et al. 2025b. Cross-continental shifts of ecological strategy in a global plant invader. Global Ecology and Biogeography 34: e70001.

Kass JM, Muscarella R, Galante PJ, Bohl CL, Pinilla-Buitrago GE, Boria RA, Soley-Guardia M, Anderson RP. 2021. ENMeval 2.0: Redesigned for customizable and reproducible modeling of species’ niches and distributions. Methods in Ecology and Evolution 12: 1602–1608.

Latzel V, Klimešová J. 2010. Transgenerational plasticity in clonal plants. Evolutionary Ecology 24: 1537–1543.

Lavoie C. 2017. The impact of invasive knotweed species (*Reynoutria* spp.) on the environment: review and research perspectives. Biological invasions 19: 2319–2337.

Leimu R, Fischer M. 2008. A meta-analysis of local adaptation in plants. PLoS One 3 : e4010.

Lowe S, Browne M, Boudjelas S, De Poorter M. 2000. 100 of the World’s Worst Invasive Alien Species: a Selection from the Global Invasive Species Database. The Invasive Species Specialist Group. Species Survival Commission of the IUCN, Auckland, 12 pp.

Li B, Suzuki J-I, Hara T. 1998. Latitudinal variation in plant size and relative growth rate in *Arabidopsis thaliana*. Oecologia 115: 293–301.

Ma H, Mo L, Crowther TW, Maynard DS, van den Hoogen J, Stocker BD, Terrer C, Zohner CM. 2021. The global distribution and environmental drivers of aboveground versus belowground plant biomass. Nature Ecology & Evolution 5: 1110–1122.

Maruta E. 1983. Growth and survival of current-year seedlings of *Polygonum cuspidatum* at the upper distribution limit on Mt. Fuji. Oecologia 60: 316–320.

McAndry C, Wilson AJ, Cotton PA, Truebano M. 2025. Transgenerational plasticity research: Challenges and opportunities. Functional Ecology 39: 3083–3104.

Mounger J, Ainouche ML, Bossdorf O, Cavé-Radet A, Li B, Parepa M, Salmon A, Yang J, Richards CL. 2021. Epigenetics and the success of invasive plants. Philosophical Transactions of the Royal Society of London. Series B, Biological Sciences, 376(1826), 20200117.

Muscarella R, Galante PJ, Soley-Guardia M, Boria RA, Kass JM, Uriarte M, Anderson RP. 2014. ENMeval: An R package for conducting spatially independent evaluations and estimating optimal model complexity for Maxentecological niche models. Methods in Ecology and Evolution 5: 1198–1205.

Naimi B, Hamm NAS, Groen TA, Skidmore AK, Toxopeus AG. 2014. Where is positional uncertainty a problem for species distribution modelling? Ecography 37: 191–203.

Oduor AMO, Leimu R, van Kleunen M. 2016. Invasive plant species are locally adapted just as frequently and at least as strongly as native plant species. The Journal of Ecology 104: 957–968.

Ohsawa H, Morita T, Shiga T, Okazaki K. 2017. Flow cytometric analysis of *Fallopia japonica* and *F*. *sachalinensis* in Japan. Journal of Phytogeography and Taxonomy 65: 7–13.

Pashley CH. 2003. The use of molecular marker in Japanese knotweed sensu lato. PhD thesis. University of Leicester. 344 pp.

Phillips SJ, Anderson RP, Schapire RE. 2006. Maximum entropy modeling of species geographic distributions. Ecological Modelling 190: 231–259.

Poorter H, Niklas KJ, Reich PB, Oleksyn J, Poot P, Mommer L. 2012. Biomass allocation to leaves, stems and roots: meta-analyses of interspecific variation and environmental control: Tansley review. The New Phytologist 193: 30–50.

Price EAC, Gamble R, Williams GG, Marshall C. 2001. Seasonal patterns of partitioning and remobilization of 14C in the invasive rhizomatous perennial Japanese knotweed (*Fallopia japonica* (Houtt.) Ronse Decraene). Evolutionary Ecology 15: 347–362.

Qi Y, Wei W, Chen C, Chen L. 2019. Plant root-shoot biomass allocation over diverse biomes: A global synthesis. Global Ecology and Conservation 18: e00606.

Radomski T. 2026. The ecology of geographic range limits. Biological Reviews of the Cambridge Philosophical Society 101: 39–59.

Richards CL, Bossdorf O, Muth NZ, Gurevitch J, Pigliucci M. 2006. Jack of all trades, master of some? On the role of phenotypic plasticity in plant invasions. Ecology Letters 9: 981–993.

R Core Team. 2025. R: A Language and Environment for Statistical Computing. R Foundation for Statistical Computing, Vienna, Austria. Available at https://www.R-project.org/.

Savolainen O, Lascoux M, Merilä J. 2013. Ecological genomics of local adaptation. Nature Reviews Genetics 14: 807–820.

Shipley BR, Bach R, Do Y, Strathearn H, McGuire JL, Dilkina B. 2022. megaSDM: integrating dispersal and time-step analyses into species distribution models. Ecography: 2022: e05450.

Simón-Porcar VI, Silva JL, Vallejo-Marín M. 2021. Rapid local adaptation in both sexual and asexual invasive populations of monkeyflowers (*Mimulus* spp.). Annals of Botany 127: 655–668.

Smith AB, Santos MJ. 2020. Testing the ability of species distribution models to infer variable importance. Ecography 43: 1801–1813.

Sommer RJ. 2020. Phenotypic plasticity: From theory and genetics to current and future challenges. Genetics 215: 1–13.

Tenkanen A, Suprun S, Oksanen E, Keinänen M, Keski-Saari S, Kontunen-Soppela S. 2021. Strategy by latitude? Higher photosynthetic capacity and root mass fraction in northern than southern silver birch (Betula pendula Roth) in uniform growing conditions. Tree Physiology 41: 974–991.

VanWallendael A, Hamann E, Franks SJ. 2018. Evidence for plasticity, but not local adaptation, in invasive Japanese knotweed (*Reynoutria japonica*) in North America. Evolutionary Ecology 32: 395–410.

Wang S, Liao Z-Y, Cao P, Schmid MW, Zhang L, Bi J, Endriss SB, Zhao Y, Parepa M, Hu W, et al. 2025. General-purpose genotypes and evolution of higher plasticity in clonality underlie knotweed invasion. The New Phytologist 246: 758–768.

Whitford AM, Shipley BR, McGuire JL. 2024. The influence of the number and distribution of background points in presence-background species distribution models. Ecological Modelling 488: 110604.

Xie X-F, Hu Y-K, Pan X, Liu F-H, Song Y-B, Dong M. 2016. Biomass allocation of stoloniferous and rhizomatous plant in response to resource availability: A phylogenetic meta-analysis. Frontiers in Plant Science 7: 603.

Younginger BS, Sirová D, Cruzan MB, Ballhorn DJ. 2017. Is biomass a reliable estimate of plant fitness? Applications in Plant Sciences 5: 1600094.

Yuan W, Pigliucci M, Richards CL. 2024. Rapid phenotypic differentiation in the iconic Japanese knotweed s.l. invading novel habitats. Scientific Reports 14: 14640.

Zhang L, van Riemsdijk I, Liu M, Lioa Z, Cavé-Radet A, Bi J, Wang S, Zhao Y, Cao P, Parepa M, Bossdorf O, Salmon A, Aïnouche M, Ju R-T, Wu J, Richards CL, Li B 2024. Biogeography of a global plant invader: from the evolutionary history to future distributions. Global Change Biology 30: e17622.

